# Su(var)3-9 mediates age-dependent increase in H3K9 methylation on TDP-43 promoter triggering neurodegeneration

**DOI:** 10.1101/2023.03.14.532519

**Authors:** Marta Marzullo, Giulia Romano, Claudia Pellacani, Federico Riccardi, Laura Ciapponi, Fabian Feiguin

## Abstract

Aging progressively modifies the physiological balance of the organism increasing susceptibility to both genetic and sporadic neurodegenerative diseases. These changes include epigenetic chromatin remodeling events that may modify gene transcription. However, how aging interconnects with disease-causing genes is not well known. Here, we found that Su(var)3-9 causes increased methylation of histone H3K9 in the promoter region of TDP-43, the most frequently altered factor in amyotrophic lateral sclerosis (ALS), affecting the mRNA and protein expression levels of this gene through epigenetic modifications in chromatin organization that appear to be conserved in aged *Drosophila* brains, mouse and human cells. Remarkably, augmented Su(var)3-9 activity causes a decrease in TDP-43 expression followed by early defects in locomotor activities. In contrast, decreasing Su(var)3-9 action promotes higher levels of TDP-43 expression and reinvigorates motility parameters in old flies, uncovering a novel role of this enzyme in regulating TDP-43 expression and locomotor senescence. The data indicate how conserved epigenetic mechanisms may link aging with neuronal diseases and suggest that Su(var)3-9 may play a role in the pathogenesis of ALS.

## Introduction

Aging is associated with a series of molecular changes, that lead to functional tissue deterioration and predispose to an increased likelihood of disease and death. This process, interestingly, does not seem to happen randomly, but follows a programmed sequence of events that appear to be conserved among evolutionarily divergent species (1–3). In the nervous system, neuronal aging or senescence can be functionally quantified through two main phenotypes, the deterioration of cognitive functions and the reduction of locomotory capacities. These alterations, on the other hand, coincide with the insidious symptoms that signal the onset and progression of the most common neurodegenerative diseases such as Alzheimer’s disease (AD), Parkinson’s disease (PD) or amyotrophic lateral sclerosis (ALS) (4,5) endorsing the idea that aging and pathological neurodegeneration may be regulated by a common set of genes (6,7).

Molecularly, a common feature of aging is the epigenetic changes in chromatin organization that occur after the post-translational modifications of histones (8,9). These modifications are conserved, affect the expression parameters of numerous genes, and may provoke alterations in the expression levels of proteins that constitute risk factors for neurodegenerative diseases. In support of this view, we and others have described that defects in the conserved TDP-43 (encoded by the *TARDBP* gene), a member of the heterogeneous nuclear ribonucleoproteins (hnRNPs) family and largely associated with the pathogenesis of ALS (10–12), is permanently required in the nervous system to maintain locomotor activity and becomes downregulated during aging in *Drosophila* and mammalian brains (13–16). Moreover, TDP-43 tight regulation in humans is required also in other tissues (such as glia and skeletal muscles) to maintain the correct molecular organization of the neuromuscular synapses and muscular innervation, all aspects critical for the motor system functioning (17–21). Even though these fluctuations in protein levels appear to be consistent and conserved in highly different species, the physiological relevance of reduced TDP-43 expression during aging, the molecules involved, and their contribution to neuronal senescence is not known. In this study, we investigated the mechanisms by which TDP-43 becomes downregulated during aging and the functional implications of these modifications in the onset and progression of locomotor waning.

## Results

### Recovery of TDP-43 function during aging prevents locomotor decline

Progressive degeneration in locomotor activity, also known as locomotor senescence, is one of the main phenotypes used to quantify the impact of age on the functional organization of the nervous system and negative geotaxis (the ability of flies to vertically climb a test cylinder) a well-accepted assay for measuring neuromuscular capacity *in vivo* (22,23). Using this methodology, we have described that the progressive decrease in *Drosophila* locomotor activity during aging correlates with a physiological decrease in the expression of the TBPH protein, homologous to the human TDP-43 (15). Consistently, we and others showed that also in mice TDP-43 undergoes an aging-dependent decrease (15,16), highlighting the evolutionary relevance of this phenomenon. However, the relationships between these events have not been clarified yet. To determine whether the drop in TBPH/TDP-43 expression during aging plays any role in locomotor senescence, we used the GeneSwitch (GS) system to generate flies carrying the neuronal driver *elav*-GS-GAL4 and the transgene UAS-TBPH (*w*^1118^; UAS-TBPH/+; *elav*-GS-GAL4/+) to modulate the expression of TBPH in a temporally controlled manner by adding the RU-486 (mifepristone) activator in the fly food (14,24,25). As controls, we utilized the TBPH^F/L^ allele unable to bind the RNA (*w*^1118^; UAS-TBPH^F/L^/+; *elav*-GS-GAL4/+) and the unrelated protein GFP (*w*^1118^; UAS-EGFP/+; *elav*-GS-GAL4/+) (26). Thus, we detected that GS-flies in which the promoter was not activated, showed a significant decrease in locomotor activity around 7 days post eclosion (dpe). This diminution in fly motility increases progressively during aging (50% at 14 dpe to 30% of their initial capacity at 21 dpe) and correlates with a decrease in TBPH/TDP-43 mRNA and protein levels (Supplementary Figure S1). Thus, to determine whether TBPH reintroduction in aged animals may prevent locomotor senescence, we induced the expression of the UAS-TBPH transgene (*w*^1118^; UAS-TBPH/+; *elav*-GS-GAL4/+) in 18 days old flies during 72 hours, by adding the RU486 activator to the fly food (Figure 1A-B). Notably, we found that induction of TBPH expression improved climbing abilities and slowed the locomotor decline in aged flies compared to UAS-TBPH^F/L^ (Figure 1C), establishing a direct correlation between the age-related decrease in TBPH expression and locomotor deterioration.

**Figure 1.**
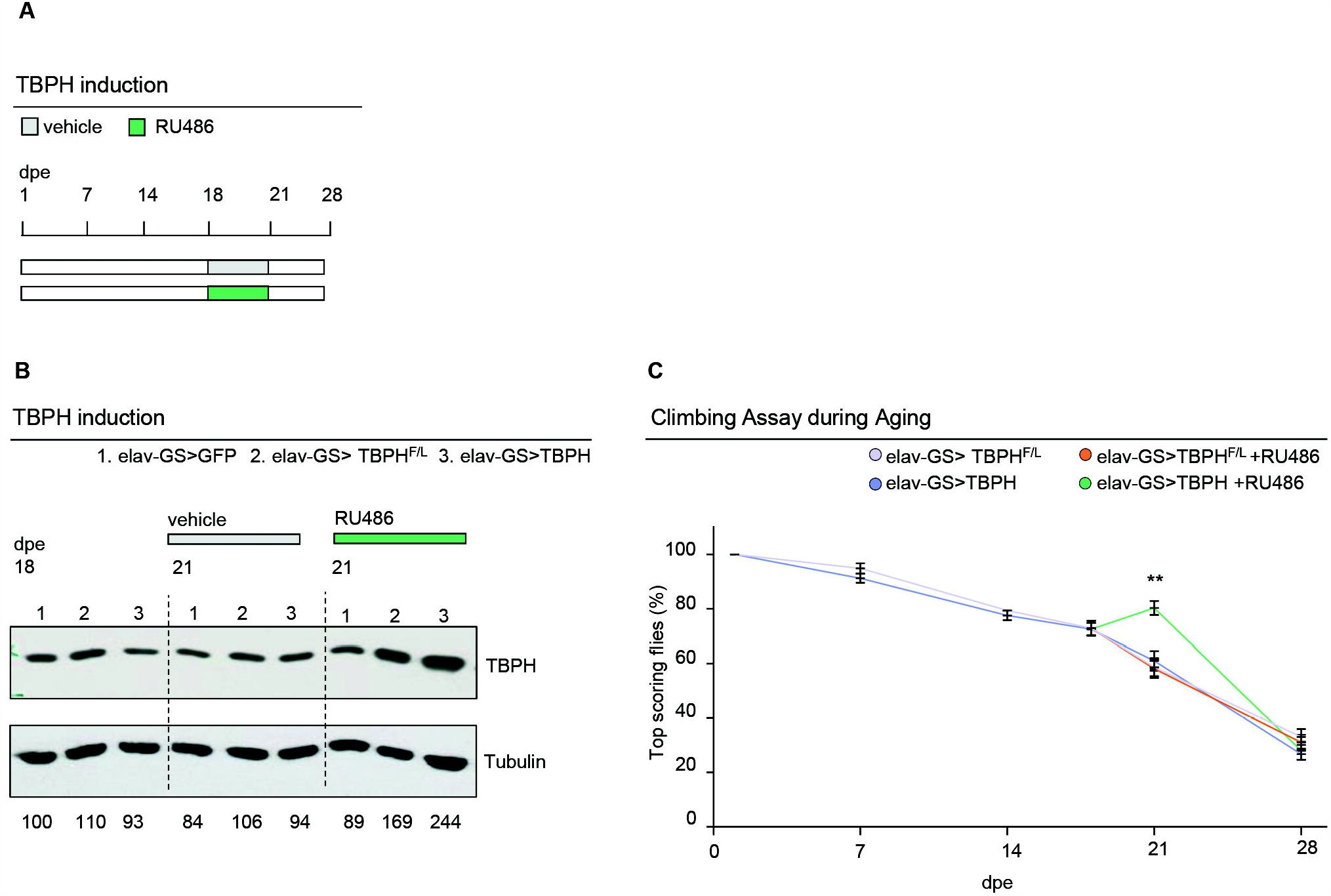
TBPH prevents locomotory senescence in Drosophila. **(A)** Schematic representation of the elav-Gene Switch induction protocol with RU486 (in green). The drug was added to fly food at day 18 until day 21, then the flies were transferred to standard food. **(B)** Western blot showing the TBPH levels in protein extracts from fly heads of the reported genotypes 1, 2 and 3 at day 18, day 21 in drug (RU486) or vehicle-only treated. Membranes were probed with anti-TBPH and anti-tubulin antibodies. Lane 1= UAS-GFPmCD8/+;*elav*GS/+; lane 2= +/+;*elav*GS/UAS-TBPH^F/L^; lane 3= UAS-TBPH/+;*elav*GS/+;. Numbers below represent band quantification normalized on internal loading (tubulin). Average of two experiments. **(C)** Climbing assay in adult flies of the reported genotypes (+/+;*elav*GS/UAS-TBPH^F/L^; and UAS-TBPH/+;*elav*GS/+), without (pink and blue, respectively) or with RU486 (orange and green, respectively) induction at different days post eclosion (7, 14, 18 and 21 dpe). Each point represents the percentage of flies able to reach the top of a 50 ml tube in 10 seconds after being tapped to the bottom. n≥ 100 animals for each genotype, in at least three technical replicates. ns, not significant; ** p<0.01 calculated by one-way ANOVA. Error bars represent SEM.

### H3K9 methylation at the *TARDBP/TBPH* promoter increases with aging and is conserved in both flies and mammals

Gene expression is a tightly regulated process influenced by the epigenetic modifications of the histones, that control the accessibility to the DNA (in particular those located in promoter regions) to a large number of proteins that can directly promote the regulation of transcription (27,28). Mechanistically, the methylation of the histone H3K9 (H3K9me) by specific methyltransferase enzymes, constitutes the initial event that triggers the formation of repressive heterochromatin domains in the DNA (29,30). Thus, to determine if the downregulation of *TBPH* during *Drosophila* aging is related to changes in the methylation patterns of H3K9, we performed chromatin immunoprecipitation (ChIP) studies and assessed the binding profile of H3K9me3 on the *TBPH* promoter. Remarkably, we found a significant enrichment in H3K9me3 amounts sited on the *TBPH* promoter in chromatin samples extracted from old flies compared to young controls (Figure 2A), revealing an increase in the levels of repressive heterochromatin modifications on the *TBPH* promoter *in vivo* during aging (16,30,31). In support of this observation, we noted that these epigenetic changes do not appear to be due to a generalized and/or nonspecific increase in H3K9 methylation caused by age, as its overall biochemical levels appear to decrease in old brains (Supplementary Figure S2), suggesting that the modifications described on the *TBPH* promoter are rather specific and may promote transcriptional repression of this gene. Importantly, we observed that similar modifications in H3K9 methylation levels of the TDP-43 promoter, take place also in the mammalian brain. Thus, H3K9me3 chromatin immunoprecipitation assays in C57 mice brains at post-natal day 10 (PND 10) and PND 365, showed a very significant increase (∼20 fold) in the methylation levels of the *TARDBP* promoter in old mice compared to young samples or to unrelated controls (Figure 2B), revealing that these modifications follow well-conserved designs.

**Figure 2.**
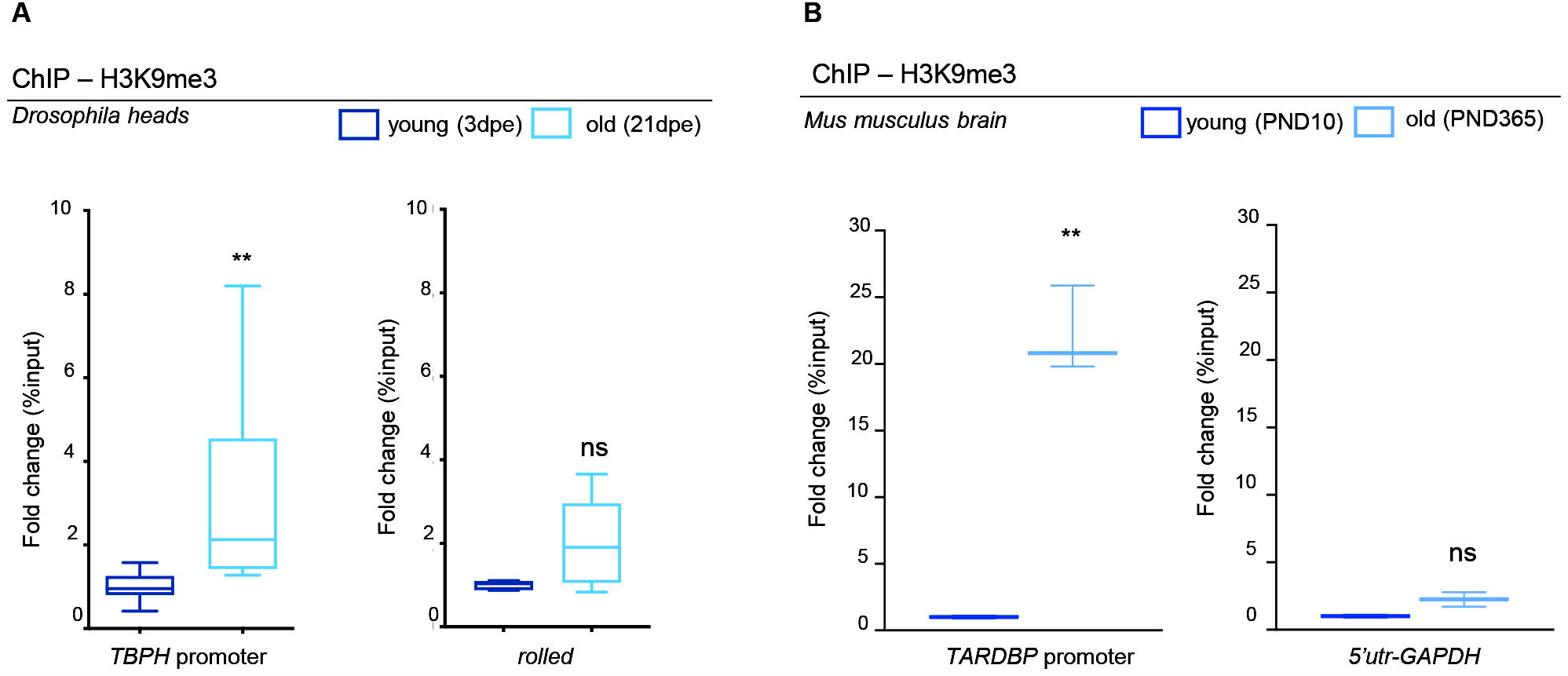
Levels of H3K9me3 at *TBPH/TARDBP* promoter increase with age. (**A**) qRT-PCR analysis on the *TBPH* promoter or on a control heterochromatic region (*rolled*), immunoprecipitated either with an anti-H3K9me3 antibody or with a control IgG antiserum in chromatin extracts from 3- or 20-days post eclosion (dpe) fly heads. The DNA enrichment is shown as a percentage of input DNA and normalized on the *GADPH* gene used as control. Note the significant increase (∼ 2 fold) of *TBPH* promoter in 21 dpe flies compared with 3 dpe. No significant changes were observed in the control gene (*rolled*). Error bars represent SEM of three independent experiments (*n* = 3; pull of 300 heads), 3 biological replicates and 3 technical replicates); **p = 0.0019, ns, not significant; Mann-Whitney t-test.(**B**) qRT-PCR analysis on the *TARDBP* promoter or on the *GADPH-5’UTR* gene used as control, immunoprecipitated either with an anti-H3K9me3 antibody or with a control IgG antiserum in the brain of C57 mice at post-natal day 10 (PND 10) or PND 365. The DNA enrichment is shown as a percentage of input DNA and normalized on the total H3. Note the significant increase (∼ 20 fold) of *m-TARDBP* promoter in PND 365 mice compared with PND 10. No significant changes were observed in the control gene *(m-GADPH*). Error bars represent SEM of three independent experiments (n = 6 mice per group, 3 biological replicates); *** p<0.001, ns, not significant; Mann-Whitney t-test.

### *Su(var)3-9* mediated H3K9 methylation of the *TBPH* promoter regulates gene expression levels and locomotor aging in flies

In order to explore the physiological significance of increased H3K9 methylation in the *TBPH* promoter region, we decided to modulate the activity of Su(var)3-9, the well-described and conserved histone methyltransferases capable of methylating H3K9 in vivo (30,32). Strikingly, we found that null alleles of *Su(var)3-9*, in trans-heterozygous combinations *(Su(var)3-9*^*6*^*/ Su(var)3-9*^*1*^), sired viable and fertile flies that present a significant increase in their locomotor capacities in adulthood compared to age-matched controls in climbing assays (Figure 3A; Supplementary video V1). Accordingly, the locomotor performance of either 20, 30, or 40 days old *Su(var)3-9* mutant flies significantly exceeded the climbing abilities of wildtype flies of the same age. Along these lines, we quantified that the loss of locomotor capacity in *Su(var)3-9* mutant flies between 3 and 30 days after hatching (from 84% of flies reaching the top to 66%, respectively) was much less pronounced than in wildtype controls (from 83% to 12%, respectively), underlining the unexpected role of this enzyme in regulating locomotor performances and neurological senescence (Figure 3A). Furthermore, biochemical analyses performed on fly head extracts obtained from the flies described above (3 and 20 days-old trans-heterozygous combinations *Su(var)3-9*^*6*^*/Su(var)3-9*^*1*^ or *w*^*1118*^ wildtype controls), revealed that both *TBPH* mRNA and protein levels are higher in *Su(var)3-9* mutants compared to the wildtype controls (Figure 3B-C). Moreover, ChIP analyses, showed that *Su(var)3-9* old mutant flies presented reduced levels of H3K9 methylation in the promoter and coding regions of *TBPH* compared to controls (Figure 3D), indicating that these molecular differences in methylation and expression levels may underlie the phenotypic changes in motility. In support of this hypothesis, we found that overexpression of *UAS-Su(var)3-9* or its human counterpart *UAS-SUV39H1*, under the control of the neuronal driver *elav-*GAL4 (Supplementary Figure S3A), was sufficient to deeply affect the locomotor capacities of these flies, inducing early locomotor decline and provoking a strong reduction in the levels of TBPH protein expression in *Drosophila* brains (Figure 3E-F), revealing that Su(var)3-9 plays a major role in the epigenetic control of TBPH expression. Additionally, we found that incubation of wildtype *Drosophila* brains with chaetocin (unfortunately the compound, in the present formulation, does not pass the gastric barrier to be tested *in vivo*) causes an increase in TBPH expression and a reduction in H3K9 methylation, mimicking the effect caused by the loss of *Su(var)3-9* (Supplementary Figure S3B). Remarkably, we observed that the role of Su(var)3-9 in the regulations of *TBPH* promoter was rather specific since the loss of two additional enzymes able to methylate H3K9, like eggless and G9a in *Drosophila* (30), was unable to modify the expression levels of *TBPH* in fly heads or affect locomotor behaviors *in vivo* (Supplementary Figure S3C-E). Curiously, we observed that the expression of the *Drosophila* homolog of Fus (dFUS-*cabeza*), a gene epistatically related to TBPH and ALS-related factor (12,33), does not change over time (Supplementary Figure S3F), suggesting that the age-dependent decline is specifically related to *TBPH* transcription.

**Figure 3.**
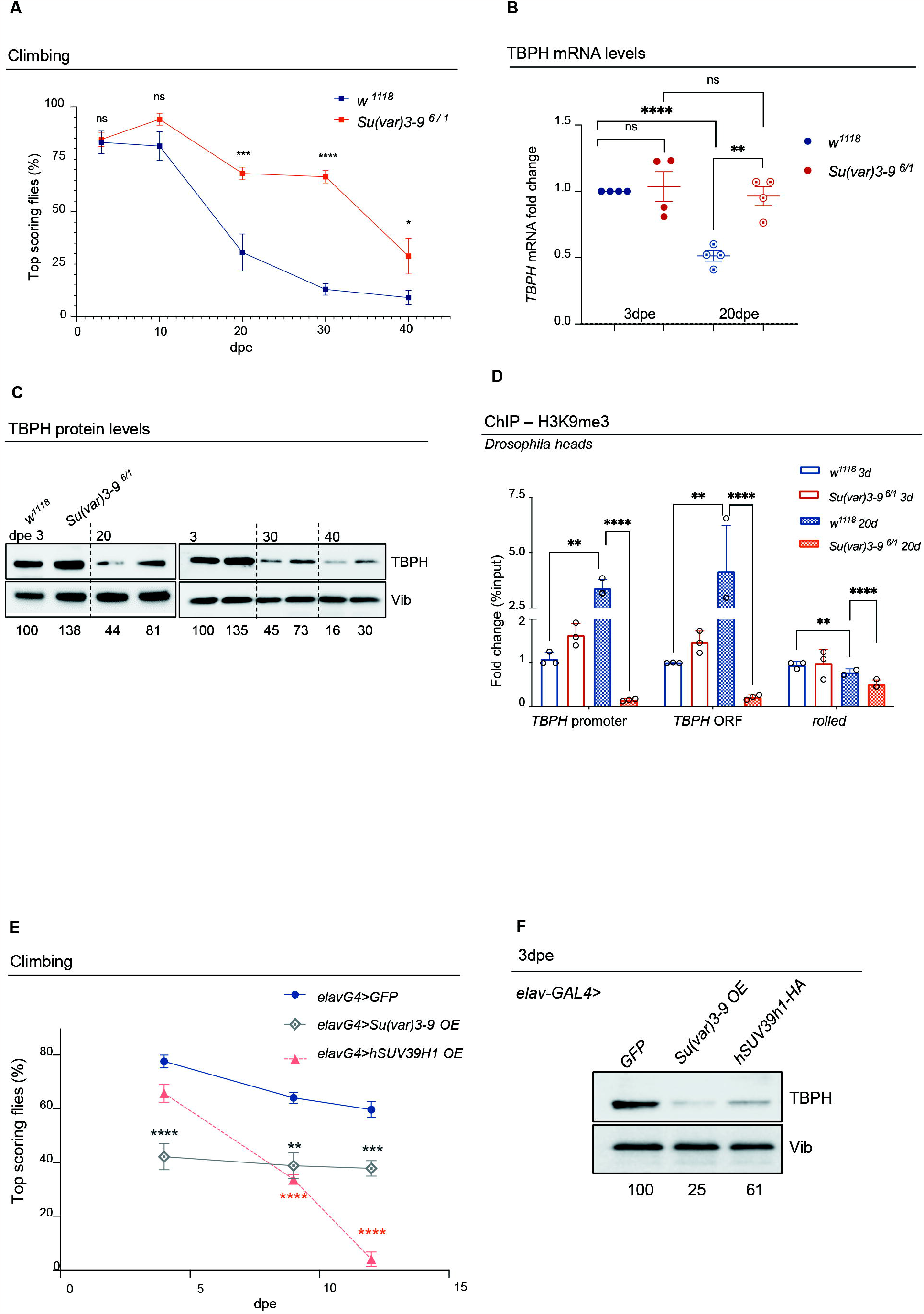
Loss of Su(var)3-9 rescues TBPH ageing-dependent decrease and associated reduced climbing abilities. (**A**) Climbing assay performed in *Su(var)3-9* mutant flies *(Su(var)3-9*^*6*^*/Su(var)3-9*^*1*^; red curve) or in control flies (*w*^*1118*^; blue curve), at different days post eclosion (3, 10, 20, 30 or 40 dpe). Each square represents the percentage of flies able to reach the top of a 50 ml tube in 10 seconds after being tapped to the bottom. n ≥30 animals for each genotype, in at least five technical replicates. ns, not significative; ***p< 0.001; ****p<0.0001 with one-way ANOVA. Error bars represent SEM. (**B**) qRT-PCR showing *TBPH* mRNA levels in *Su(var)3-9* mutants [*Su(var)3-9*^*6*^*/Su(var)3-9*^*1*^; red] compared to controls (*w*^*1118*^; blue) in RNAs from young (3dpe; full circles) or old (20 dpe; empty-dotted circles) flies heads extracts. Error bars represent SEM of three independent experiments (*n* = 3; pull of 50 heads), 3 biological replicates and 3 technical replicates). ** p<0.01; ****p<0.0001 with one-way ANOVA. (**C**) Western Blot showing the TBPH protein levels in *Su(var)3-9* mutants [*Su(var)3-9*^*6*^*/Su(var)3-9*^*1*^] compared to controls (*w*^*1118*^) in fly head extracts at different days post eclosion (3, 20, 30 or 40 dpe). Numbers below represent band quantification (the average of four experiments) normalized on internal loading (Vibrator, Vib). (**D**) qRT-PCR analysis on both the *TBPH* promoter and its coding sequence compared to a control heterochromatic region (*rolled*), immunoprecipitated either with an anti-H3K9me3 antibody or with a control IgG antisera in chromatin extracts from young (3 dpe) or old (20 dpe) *Su(var)3-9* mutants [*Su(var)3-9*^*6*^*/Su(var)3-9*^*1*^] or controls (*w*^*1118*^). The DNA enrichment is shown as a percentage of input DNA and normalized on the *GAPDH* gene used as control. Error bars represent SEM of three independent experiments (*n* =3; pull of 300 heads, 3 biological replicates and 3 technical replicates). ** p<0.01; ****p<0.0001 with one-way ANOVA. (E) Climbing assay performed in adult flies overexpressing *UAS-Su(var)3-9* (gray curve), or *UAS-hSuv39h1-HA* (*orange curve*) under the control of the *elav-GAL4* driver or in control flies expressing a *UAS-GFP* construct (blue curve), at different days post eclosion (3, 7, or 12 dpe), at 29°C. Each dot represents the percentage of flies that reach the top of a 50 ml tube in 10 seconds after being tapped to the bottom. n≥ 30 animals for each genotype, at least 5 technical replicates. ** p<0.01 *** p<0.001; ****p<0.0001 calculated by one-way ANOVA. (F) Western Blot showing the TBPH protein levels in heads extracts of flies overexpressing the *UAS-Su(var)3-9* or the *UAS-hSuv39h1-HA* or *UAS-GFP* under the control of the *elav-GAL4* driver at 3 days post eclosion. Numbers below represent band quantification normalized on internal loading (Vibrator, Vib).

### The conserved SUV39H1 enzyme regulates TDP-43 expression levels in human cells

To determine if the conserved SUV39H1 histone methyltransferase is able to regulate the methylation of the TDP-43 promoter and modulate protein expression also in human cells, we took advantage of a HaCaT cell line carrying a CRISPR-Cas9 mutation in the *SUV39H1* gene (*SUV39H1 KO;* (34). Interestingly, we observed that in these cells the absence of SUV39H1 causes an increase in the levels of TDP-43 protein expression (Figure 4A). In the same direction, H3K9me3 ChIP analyses revealed a significant reduction in H3K9me3 amounts sited on the *TARDBP* promoter in chromatin samples extracted from SUV39H1 KO cells compared to wildtype cells (Figure 4B), suggesting that epigenetic modifications mediated by SUV39H1 might be responsible for the transcriptional repression of *TARDBP* and, above all, underlining the remarkable conservation found in the regulation of this locus. To challenge whether aging-induced modifications would also play a role in the regulation of human TDP-43, we treated wildtype or SUV39H1 KO cells with H_2_O_2_ a classic and well-accepted treatment for inducing cellular senescence (35–37); (Supplementary Figure S4). As a result, we found that H_2_O_2_ induced a significant reduction in TDP-43 protein expression which is prevented by the deletion of the *SUV39H1* gene (Figure 4C), indicating that similar age-dependent regulatory mechanisms might be present in human cells.

**Figure 4.**
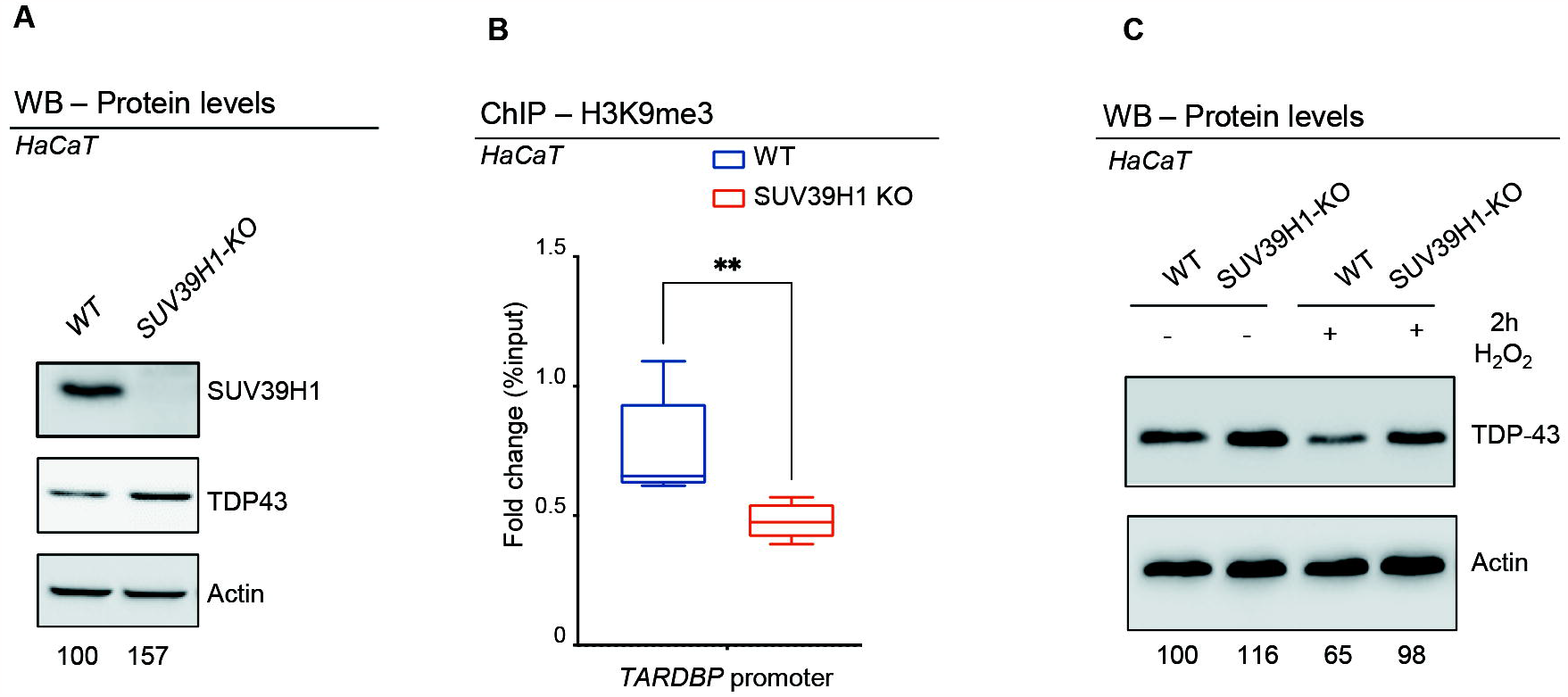
SUV39H1 depletion in human cells correlates with reduced levels of H3K9me3 at *TARDBP* promoter and with a corresponding increase in TDP-43 protein. (**A**) Western Blot showing the SUV39H1 and TDP-43 protein levels in extracts from WT or SUV39H1 KO cells. Numbers below represent band quantification normalized on internal loading control (actin; average of 6 experiments). (**B**) qRT-PCR analysis on the *hTARDBP* promoter immunoprecipitated with an anti-H3K9me3 in chromatin extracts from WT or SUV39H1 KO cells. Enrichment is shown as a percentage of input DNA and normalized on the *GADPH* gene used as control. Error bars represent SEM of three independent experiments (n = 3, 3 biological replicates). (**C**) Western blots showing the expression levels of TDP-43 in wild type (WT) or *SUV39H1 KO* HaCaT Keratinocytes after (+) or not (−) treatment with H_2_O_2_ (200mM) for 2 ours (2h). H_2_O_2_ treatment reduces TDP-43 levels in WT but not in SUV39H1-KO cells. Numbers below represent band quantification normalized on internal loading control (actin; average of 3 biological repetitions).

## Discussion

One of the most fundamental features of aging is the progressive deterioration in locomotor skills. Despite some studies, in both mice and flies revealing that TDP-43/TBPH levels decrease during aging (15,16,38), the mechanism underlying aging-dependent locomotor senescence and the accompanying physiological decrease of TDP-43 is unclear. Likewise, it is not obvious whether the lowering of TDP-43 levels leads to a locomotor decline in the elderly. In this manuscript, we found that induction of the TDP-43 fly counterpart, TBPH, expression in old fly neurons, but not of the TBPH^F/L^ mutated form (unable to bind RNA), is sufficient to rescue locomotor senescence, demonstrating a direct correlation between these events and revealing a novel role for TDP-43/TBPH in the regulation of age-dependent locomotor degeneration. In that direction, alteration in the function of TDP-43 is considered one of the main causes of ALS and it has been shown that pathological variations in the intracellular levels of this protein (both gain or loss of function) were able to cause neuronal death indicating that tight control of TDP-43 expression is crucial to prevent neurological phenotypes (39–41). These observations, therefore, highlight the importance that the knowledge of novel genes or molecules capable to modulate TDP-43 activity could have for understanding the pathogenesis of ALS (42–45). According to that, we found an age-dependent increase in H3K9 methylation at the TBPH/TDP-43 promoter region mediated by *Su(var)3-9* in *Drosophila* and confirmed that these modifications are conserved in mice brains and human cells. Moreover, we established that these regulatory mechanisms were sufficient to modulate the expression levels of TDP-43 in both flies and human cells and to affect locomotor behaviors. Interestingly, a similar outcome was detected using chaetocin, a chemical compound capable of inhibiting Su(var)3-9-mediated H3K9 methylation (46). These data reinforce the idea that Su(var)3-9 plays a fundamental role in the epigenetic regulation of *TBPH* expression and identifies a compound capable of regulating the expression levels of this gene *in situ*, contributing to the development of potential pharmacological interventions against ALS or locomotor weakening in the future.

In connection with the mechanisms involved, it is unclear how aging might influence the activity of Su(var)3-9 or the methylation status of the TBPH/TDP-43 promoter considering that the total amount of H3K9 methylation in the genome decreases with age (Supplementary Figure S2) (47). Intriguingly, we found that the presence of aging-related factors able to induce early senescence in cultured human cells, such as H_2_0_2_ accumulation, leads to a reduction in TDP-43 protein expression mediated by the human-homolog gene *SUV39H1* (Figure 4C), suggesting that the metabolic changes that precede and drive aging could modify/increase the function of this enzyme in specific regions or loci of the chromosome (48,49). In agreement with this idea, ChIP array analyses performed in *Drosophila* brains found the increased accumulation of the *Su(var)3-9* protein itself at the promoter region of TBPH during aging (50,51). In addition, we detected a slight but significant increase in *Su(var)3-9* expression in old flies (Supplementary Figure S3G), explaining how the accumulation of epigenetic modifications in this locus could happen over time (Figure 5). In any case, additional experiments are necessary to deepen our knowledge of the issues discussed above.

**Figure 5.**
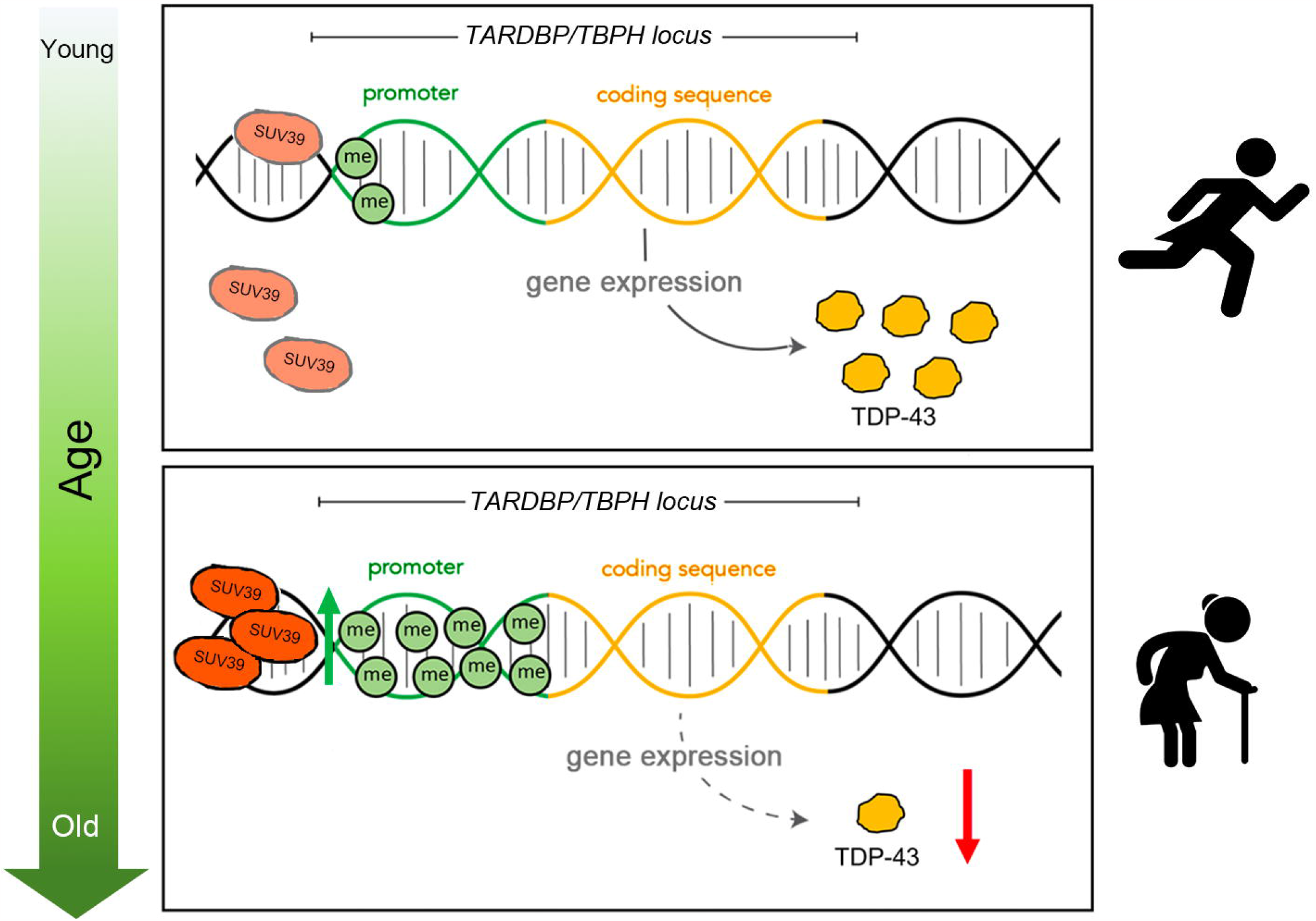
Schematic representation of the mechanism of action of Suv39 at *TARDBP/TBPH* promoter region during aging. SUV39 activity at the *TARDBP/TBPH* promoter region is increased in elderly individuals. This effect results in increased methylation of H3K9 leading to reduced levels of TDP-43 expression and diminished locomotor capabilities.

In conclusion, we have identified an unprecedented mechanism whereby Su(var)3-9 regulates the epigenetic status of the *TARDBP*/*TBPH* promoter and drives the progression of locomotor aging through the regulation of TDP-43/*TBPH* expression levels. This role of SUV39 seems to be evolutionarily conserved from *Drosophila* to vertebrates and may help to understand the interrelationships between human aging and neurodegenerative diseases.

## Acknowledgments

We thank Dr. Wiesława Leśniak (Nencki Institute of Experimental Biology, Warsaw, Poland) for the SUV39H1 mutant cell lines. Prof. Gunter Reuter (Institute of Biology and Developmental Genetics, Halle, Germany), Dr. Marion Delattre (Department of Genetics and Evolution, University of Geneva, Geneva, Switzerland) for the *Su(var)3-9*^*6*^ and *G9a*^*RG5*^ fly stocks respectively, and Prof. Franco Pagani (ICGEB-Trieste) for sharing reagents. This work was supported by AFM-Telethon (project 21025) and AriSLA (NOSRESCUEALS) awards to L.C., and F.F.

## Author contributions

M.M., G.R., L.C., and F.F. designed the experiments. M.M., G.R., C.P., and F.R. performed the experiments and collected the data, and analyzed the results together with L.C., and F.F. M.M., G.R., L.C., and F.F. wrote the manuscript.

## Methods

### Drosophila strains and rearing conditions

Drosophila stocks were maintained on standard fly food (25 g/L corn flour, 5 g/L lyophilized agar, 50 g/L sugar, 50 g/L fresh yeast, 2,5 mL/L Tegosept [10% in ethanol], and 2.5 mL/L propionic acid) at 25 □C in a 12h light/dark cycle. All experiments were performed in the same standard conditions, otherwise differently specified. The following fly strains were purchased from the Bloomington Drosophila Stock Center (BDSC, Indiana University, Bloomington, IN, USA): *w*^*1118*^ (BDSC #3605); *elav*-GS (BDSC #43642); *UAS-TBPH* (BDSC #93601); *UAS-TBPH*.*F-L* (BDSC #93781); *UAS-mCD8-GFP* (BDSC #30002); *Su(var)3-9*^*1*^/TM3 (BDSC #6209); UAS-Su(var)3-9.lacI (BDSC #93147); *UAS-hSUV39H1*.*HA* (BDSC #84799); *elav-GAL4* (BDSC #77894); *egg*^1473^/SM1 (BDSC #30565). The *Su(var)3-9*^*6*^/TM6B allele was a kind gift of Gunter Reuter (52), the *G9a*^*RG5*^ allele was a kind gift of Marion Delattre (53).

### Climbing assays

The locomotion activity was measured by quantification of geotactic response. Equal ratio of male and females of the desired ages will be transferred, without anesthesia, to a 15 ml conical tube, tapped to the bottom of the tube, and their subsequent climbing activity quantified as the percentage of flies reaching the top of the tube in 10s (54). The number of climbing events was scored for 5 consecutive times. Flies were assessed in batches of 15, at least three biological replicates were performed for each condition (13).

### RU486-Induction protocol

The Gene Switch system was activated by adding the RU486 (Sigma-Aldrich #M8046) activator to the fly food. A stock solution of 50 mM RU486 in 95% ethanol was diluted to the final concentration of 0.5 mM in 2% sucrose and the solution was been added on the surface of standard cornmeal medium to feed adults.

### Chaetocin treatment

Adult fly heads were separated from the bodies and incubated with 100 nM chaetocin (Sigma-Aldrich #C9492) or 100% Ethanol in Schneider’s Medium supplemented with 10% FBS for 2 h at room temperature. Heads were then washed in PBS1x and collected for subsequent analysis.

### Chromatin immunoprecipitation

#### Fly heads

Heads of frozen flies were separated by vortexing for 15 sec and isolated using 630 μm and 400 μm sieves. 400 – 600 fly heads were homogenized in homogenization buffer [350 mM sucrose, 15 mM HEPES pH 7.6, 10 mM KCl, 5 mM MgCl2, 0.5 mM EGTA, 0.1 mM EDTA, 0.1% Tween, with 1 mM DTT and Protease Inhibitor Cocktail (PIC, Roche) added immediately prior to use] at 4 °C. The homogenate was fixed using 1% formaldehyde for 10 min at RT and then quenched with glycine. The tissue debris was removed by filtration with 60 μm nylon net (Millipore). Nuclei were collected and washed with RIPA buffer at 4 °C (150 mM NaCl, 25 mM HEPES pH 7.6, 1 mM EDTA, 1% Triton-X, 0.1% SDS, 0.1% DOC, with protease inhibitors added prior to use). The extract was sonicated 6 times with 2 min cycles (Branson Sonifier 250, output=50%). Sonicated samples were centrifuged for 10 min at 12,000 × g. Two hundred and fifty micrograms of chromatin DNA were subjected to a 1 h preclearing with 50 *μ*l of a 50% protein G-Sepharose (GE healthcare) bead slurry containing 1% BSA. Before the Immunoprecipitation, 5% of the total extract was collected as INPUT. The precleared samples were then immunoprecipitated overnight with 5 *μ*g of anti-H3K9me3 (Abcam ab8898) or anti-rabbit IgGs (Sigma, 15006) at 4 °C. The immune complexes were incubated for 4 h at 4 °C with 50 *μ*l of fresh protein G-Sepharose beads. After immunopurification, beads were washed four times with RIPA and once with LiCl wash buffer (250 mM LiCl, 10 mM Tris-Hcl pH 8.0, 1 mM EDTA, 0.5% NP-40, protease inhibitors PIC (Roche). Beads were re-suspended in TE buffer and incubated ON at 65°C. Proteins were digested with Proteinase K (10 mg/ml) at 55 °C for 1 h. Immunoprecipitated DNA was purified using Phenol:Chlorophorm:Isoamyl alchol extraction. Immunoprecipitated DNA and 5% input DNA were analyzed by SYBR-Green real-time qPCR. The run was performed by using the Applied Biosystems (Waltham, MA) Quant-Studio 3 Real-Time PCR System 36 instrument. Primer Sequences described previously are reported in Table S1.

### Mouse brain

Chromatin immunoprecipitation in brain of C57 mice at post-natal day 10 (PND 10) and PND 365 was performed using EpiQuik Tissue Chromatin Immunoprecipitation kit (Epigentek #P-2003) according to manufacturer’s instructions. Briefly, 150 mg of frozen tissue were cut into small pieces (<1 mm^3^) and cross-linked with 1% formaldehyde for 10 min at room temperature and then quenched in PBS 1X-Glycine 1.25M for 10 min at room temperature. The samples were homogenized using a Douncer homogenizer and centrifuged to pellet nuclei. After homogenization, lysis buffer was added to nuclei. Chromatin was prepared and sonicated using a water bath Bioruptor (Diagenode; 30” ON/30” OFF, High power, 3 × 10 cycles) to a size range of 200 –1000 bp. To pre-cleared cell debris, sonicated chromatin was centrifuged at 12,000 × g at +4°C for 10 minutes. Chromatin was diluted and ChIP performed according to manufacturer’s instructions using antibodies against H3K9me3 (ab8898, Abcam), histone H3 (ab1791, Abcam), IgG1 (G3A1, Cell Signalling) was used as negative control in the immunoprecipitation. Immunoprecipitated DNA was purified by phenol-chloroform extraction and in parallel 5 ul (5%) were taken to be used as input in the quantification analysis. qPCRs were performed using iQ SYBR Green in a CFX96 Real-Time PCR system (Bio-Rad). Primer sequences are reported in Table S1.

### Human HaCaT cells

HaCaT cells were crosslinked with 1% formaldehyde fixing buffer (1% Formaldehyde; 5 mM Hepes pH8.0; 0.05 mM EGTA pH 8.0; 10 mM NaCl) at 37 °C for 10 minutes and then quenched with glycine, rinsed twice with cold phosphate-buffered saline, and then lysed and harvested in ChIP lysis buffer (50 mM Tris-HCl pH 8.1; 0.5% SDS; 10 mm EDTA;100 mM NaCl, 1mM PMSF, Proteinase inhibitor Roche) by centrifugation for 6 min at 2,000 × *g*. Cells were then resuspended in sonication buffer (50 mM Tris-HCl pH 8.1; 10 mm EDTA; 1% Triton-X; 0,1% deoxycholate sodium; 100 mM NaCl, 1mM PMSF, Proteinase inhibitor Roche) and sonicated 6 times with 2 min cycles (Branson Sonifier 250, output=50%). Sonicated samples were centrifuged for 10 min at 12,000 × *g* and the supernatant were diluted 5-fold in sonication buffer. Two hundred and fifty micrograms of chromatin DNA were subjected to a 1 h preclearing with 50 *μ*l of a 50% protein G-Sepharose (GE healthcare) bead slurry containing 1% BSA. Before the Immunoprecipitation, 5% of the total extract was collected as INPUT. The precleared samples were then immunoprecipitated overnight with 5 *μ*g of anti-H3K9me3 (Abcam ab8898) at 4 °C. The immune complexes were then incubated for 4 h at 4 °C with 50 *μ*l of fresh protein G-Sepharose beads. Following incubation, the beads were collected by centrifugation for 1 min at 2,000 × *g* and washed consecutively for 3–5 min with 1 ml of each solution: low-salt wash buffer (0.1% SDS; 1% Triton X-100; 2 mM EDTA; 20 mM Tris pH 8.1; and 150 mM NaCl), high-salt wash buffer (0.1% SDS; 1% Triton X-100; 2 mM EDTA, 20 mM Tris pH 8.1; and 500 mM NaCl), LiCl wash buffer (250 mM LiCl; 1% NP-40, 1% deoxycholate sodium salt, 1 mM EDTA, and 10 mM Tris pH 8.1), and twice in Tris and EDTA buffers (10 mM Tris pH 8.1 and 1 mM EDTA). Immune complexes were then eluted with 120 *μ*l of buffer containing 1% SDS and 100 mm NaHCO_3_. Crosslinking was reversed by incubating the samples overnight at 65 °C. Proteins were digested with Proteinase K (10 mg/ml) at 55 °C for 1 h. Immunoprecipitated DNA was purified using Phenol:Chlorophorm:Isoamyl alcohol extraction. Immunoprecipitated DNA (1.5 *μ*l) and 5% input DNA were analyzed by SYBR-Green real-time qPCR (as described in Antonucci et al 2014). The run was performed by using the Applied Biosystems (Waltham, MA) Quant-Studio 3 Real-Time PCR System 36 instrument. Primer Sequences described previously are reported in Table S1.

### RNA extraction and quantitative PCR

Total mRNA was isolated from Drosophila adult heads by using Trizol reagent (15596026, Thermo Fisher Scientific) according to the manufacturer’s instructions. RNA was reverse-transcribed (1 mg each experimental point) by using SensiFAST cDNA Synthesis Kit (BIO-65053, Bioline) and qPCR was performed as described (18) using SensiFast Sybr Lo-Rox Mix (BIO-94020, Bioline). The run was performed by using the Applied Biosystems (Waltham, MA) Quant Studio 3 Real-Time PCR System 36 instrument. Primer Sequences are reported in Table <u>S1</u>.

### Human HaCAT cells

The immortalized human epidermal keratinocyte (HaCaT) cell line was obtained from (19). The HaCaT cells were cultured in complete media, which comprised of Dulbecco’s modified Eagle’s medium (DMEM) supplemented with 10% (v/v) heat-inactivated fetal bovine serum and 1% (v/v) penicillin-streptomycin at 37 °C in a humidified atmosphere of 5% CO_2_/95% air.

### H_2_O_2_ Treatment

HaCaT cells (10^5^cells) were cultured on 35 mm cell culture dish for 24 h and treated with H_2_O_2_ at 200 *μ*M/l for 2 h at 37 °C.

H_2_O_2_ was washed with PBS for terminating the treatment. Cells were kept on the incubation in normal medium for another 24 h. Cells were then harvested and assessed in western blot.

### Western Blot

#### Fly extract

Protein extracts were derived from adult fly heads, lysed in sample buffer or Urea Buffer (150 mM NaCl, 10 mM Tris-HCl pH8, 0,5 mM EDTA, 10% glycerol, 5 mM EGTA, 50 mM NaF, 4 M urea, 5 mM DTT, Protease Inhibitor Cocktail (PIC) (Roche), fractionated by SDS-PAGE and transferred to nitrocellulose membrane. Primary antibodies were: anti-TBPH rabbit (1:1000; homemade (13); anti-Actin goat (1:1000; Santa Cruz, sc-1616); anti-Vibrator rabbit (1:5000; also named Giotto (55); anti-H3K9me2 mouse (1:400; Abcam ab1220), anti-H3K9me3 rabbit (1:1000; Abcam ab8898); anti-Tubulin mouse (1:5000; Sigma, T-5168); anti-HA HRP (1:1000; Santa Cruz sc7392); anti-Su(var)3-9 rat (1:50; (32). As a secondary antibody, we used the appropriate HRP-conjugated antibody (GE Healthcare) diluted 1:5000 in PBS-Tween 0.1%. Membranes were incubated 5□min with ECL substrate (#1705062 and #1705060, Bio-Rad) and the HRP-ECL reaction was revealed using the ChemiDoc™ XRS gel imaging system (Bio-Rad). Band intensity quantification was performed using the gel analyzer tool in Fiji/ImageJ software.

### HaCAT extract

Cells were harvested and centrifuged at 5,000 rpm for 5 min at 4°C. The supernatant was removed, Buffer WCE 2X (100 mM TrisHCl pH 6.8, 4% SDS, 200 mM DTT) was added to resuspend the cell pellet, boiled for 10 min and then added an equal volume of SDS-PAGE Sample Loading Buffer [2X] (100 mM TrisHCl pH 6.8, 4% SDS, 200 mM DTT, 20% glycerol, 0.004% bromphenol blue) to the mixture. Cell extracts were pelleted at 15,000 g in an Eppendorf centrifuge for 15 min at 4°C and the supernatants were analyzed by Western blotting according to (56), using the following antibodies, all diluted in TBS-T: anti-p-p53 (Ser 15; 1:1000, Santa Cell Signaling), anti-p53 (1:1000, Santa Cruz), anti-p-H2AX (Ser 139; 1:1000, Millipore), anti-SUV39H1 (44.1; 1:1000, Santa Cruz Biotechnology), anti-TDP-43 (1:5000, Proteintech), anti-H3k9me3 (1:1000, Abcam ab8898), anti-H3K9me2 (1:500, Abcam ab1220), anti-actin-HRP-conjugated (1:5000, Santa Cruz Biotechnology). These primary antibodies were detected using HRP conjugated anti-mouse and anti-rabbit IgGs and the ECL detection kit (all from GE Healthcare). Band intensities were quantified by densitometric analysis with Image Lab software (Bio-Rad).

All full length uncropped original western blots are available in the Supplementary Materials section.

### Statistical analyses

Statistical analysis was performed using Prism six software (MacKiev). The Shapiro-Wilk test was used to assess the normal distribution of every group of different genotypes. Statistical differences for multiple comparisons were analyzed with the Kruskal-Wallis for non-parametric values or with one-way ANOVA for parametric values. The Dunn’s or the Tukey’s test was performed, respectively, as post hoc test to determine the significance between every single group. The Mann-Whitney U-test or the t-test were used for two groups’ comparison of non-parametric or parametric values, respectively. A p< 0.05 was considered significant.

