## Supplementary Material for "Su(var)3-9 mediates age-dependent increase in H3K9 methylation on TDP-43 promoter triggering neurodegeneration"

**Table S1. List of primer used in this work:**

| ***Name*** | **Sequence** | **Reference** |
|---|---|---|
| *TBPH_real-time FW* | CGGCAAGCCGAGCACGATGAG | (1) |
| *TBPH_real-time RV* | CGCGGAGTTCGCTCCAACGAG | |
| *TBPH_promoter FW* | CACTAGCCCTGTTCGGCCAC | This study |
| *TBPH_promoter RV* | AGCATTTCTTCTCGCTTTCCGTT | |
| *rolled_FW* | TTGGATTGGCTCGTATTGCA | This study |
| *rolled_RV* | CCATCGGGTAGCAACGTATTC | |
| *GAPDH_FW* | CCTGGCCAAGGTCATCAATG | (2) |
| *GAPDH_RV* | ATGACCTTGCCCACAGCCTT | |
| *TDP-43_promoter Fw (mouse)* | GAAGCCAGTGGGAGAGG | This study |
| *TDP-43_promoter RV (mouse)* | ACAACAGCCGCGCTACC | |
| *GAPDH-5'UTR FW (mouse)* | GGGTTCCTATAAATACGGACTGC | |
| *GAPDH-5'UTR RE(mouse)* | CTGGCACTGCACAAGAAGA | |
| *caz(dFus)_FW* | GCAATTTGTACGCAGCGAGT | This study |
| *caz(dFus)_RV* | TGGCTCGTTGTAGTTACCCG | |
| *human_TARDBP_promoter FW* | CAACCGGTGGGAGAGGACGCCG | This study |
| *human_TARDBP_promoter FW* | CGCGACTCACCCGCTAGGCCG | |
| Su(var)3-9_FW | CCACGGTGGTCAAAGCCATA | This study |
| Su(var)3-9_RV | GGCTATGTCGCCGCAATTC | |
| *TBPH_ORF FW* | TTCCCAAGGGTAAATTCGAG | This study |
| *TBPH_ORF RV* | TGGCAGTTGAGTTCTCCAAG | |

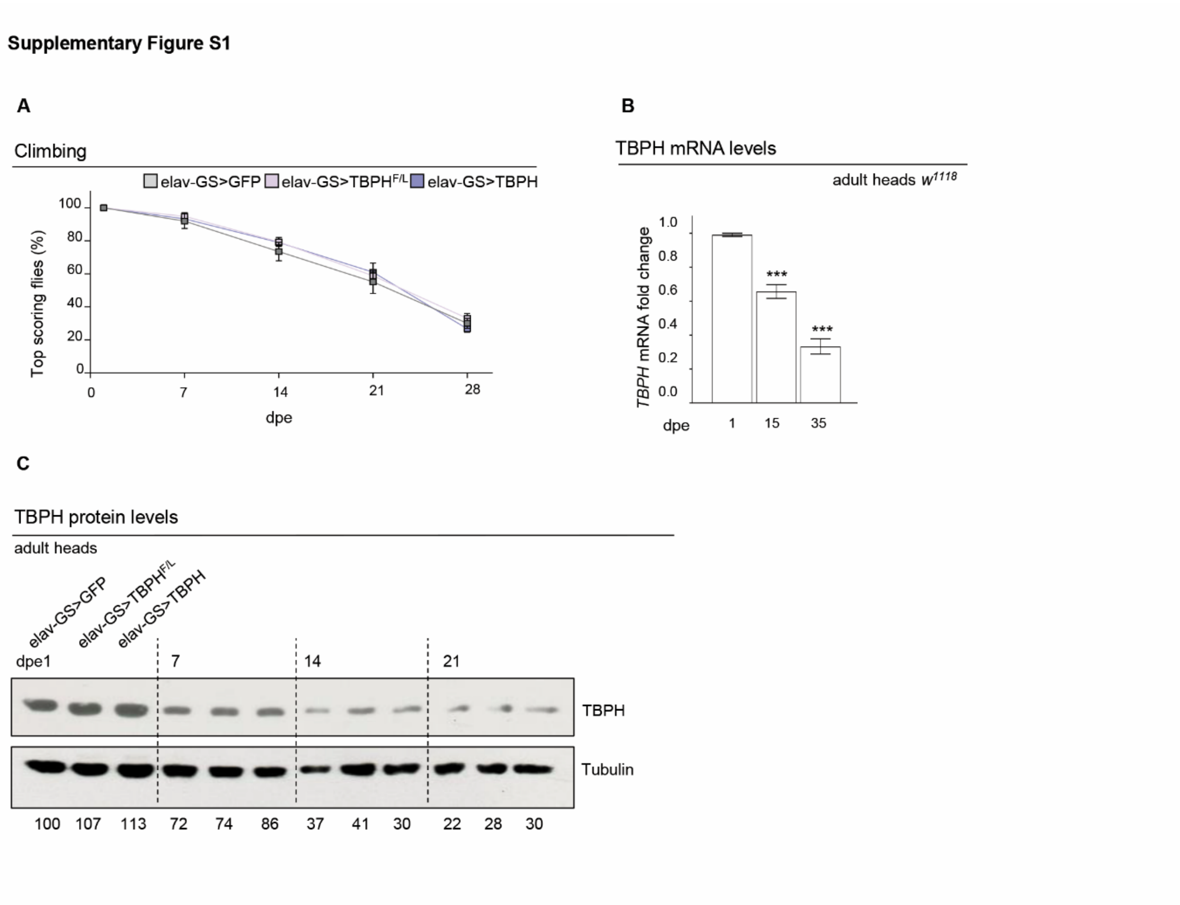


**Supplementary Figure S1. Locomotor decline correlates with reduction of TBPH levels in Drosophila**

(**A**) Climbing assay performed on adult flies of the reported genotypes at different days post eclosion (7, 14, 21, or 28 dpe) without induction with RU486 of the *elav*-GeneSwitch GAL4 driver. Each box represents the percentage of flies able to reach the top of a 50 ml tube in 10 seconds after being tapped to the bottom. n≥ 80 animals for each genotype, in at least three technical replicates. ns, not significant with one-way ANOVA. Error bars represent SEM. (**B**) qRT-PCR showing *TBPH* mRNA levels in RNAs from wild type ($w^{1118}$) fly head extracts. Error bars represent SEM of 3 biological replicates. *** $p<0.001$, by one-way ANOVA. (**C**) Western Blot showing the TBPH levels in wild type ($w^{1118}$) protein extracts from fly heads of *elav*-GeneSwitch flies driving UAS-GFP (1), UAS-$TBPH^{F/L}$ (2) or UAS-TBPH (3) without RU486 induction at different days post eclosion (7, 14 and 21 dpe). Numbers below represent band quantification normalized on internal loading (tubulin).

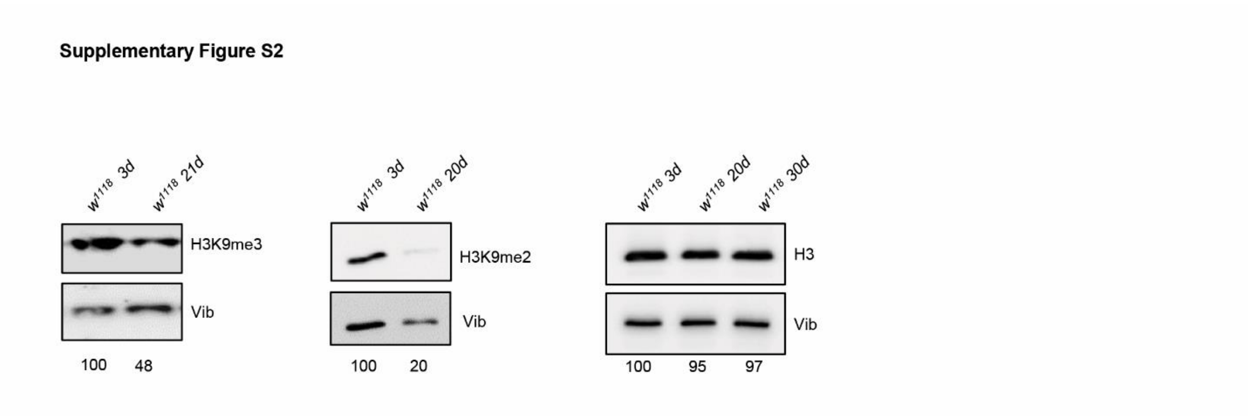


**Supplementary Figure S2. Global H3K9 methylation decreases during aging.**
Western Blot on protein extracts from adult fly heads at 3 or 20dpe detected with anti-H3K9me3, anti-H3K9me2 and anti-H3 showing that the global levels of H3K9 methylation undergo an aging-dependent reduction, while the total histone H3 does not change. Numbers below represent band quantification normalized on internal loading (Vibrator, Vib). Average of two experiments.

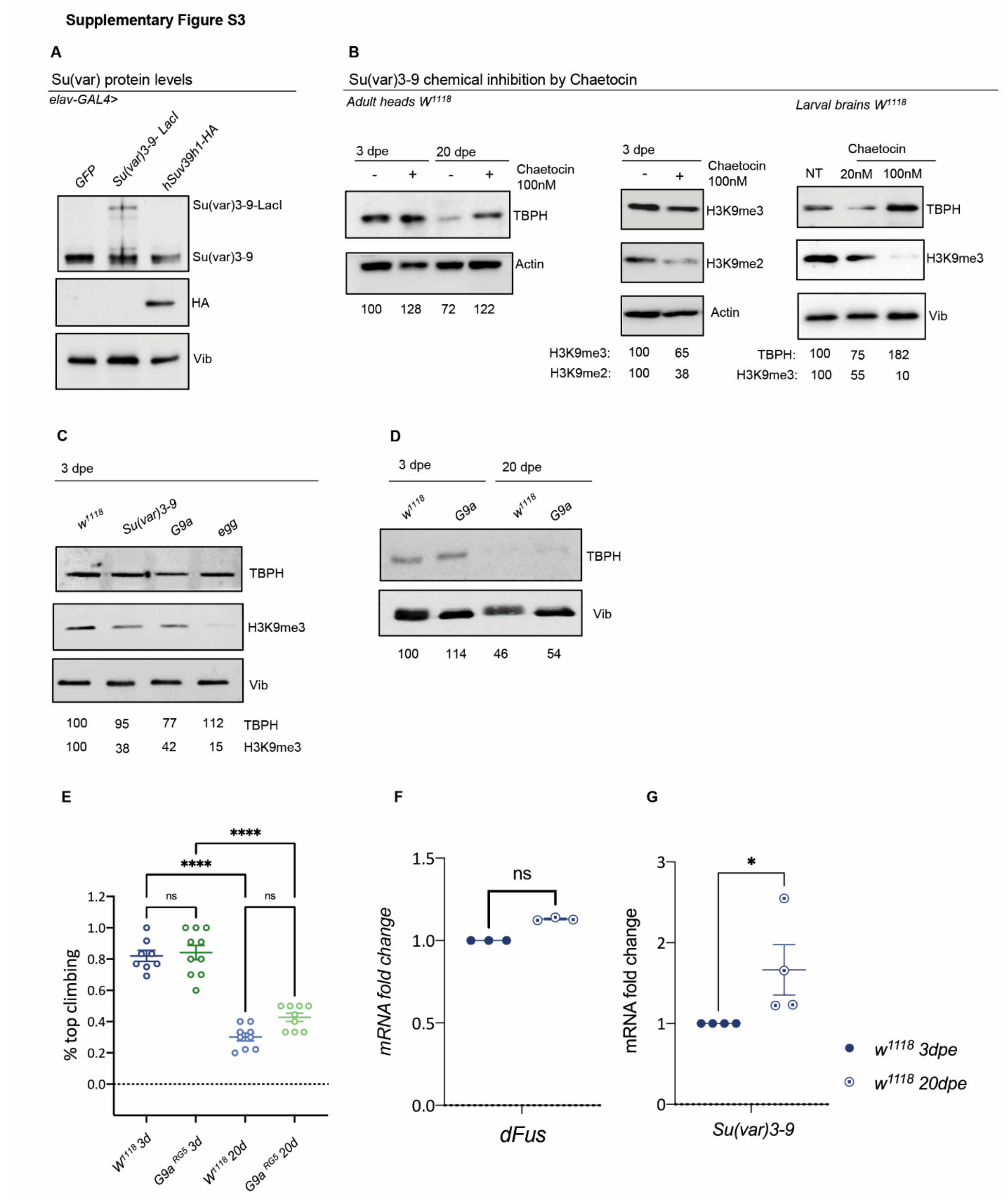


**Supplementary Figure S3. Su(var)3-9 specifically controls TBPH expression levels.**
(**A**) Western Blot showing the SUV3-9 proteins level in heads extracts of flies overexpressing either the UAS-*Su(var)3-9-lacI*, the *UAS-hSUV39h1-HA* or *UAS-GFP* construct under the control

of the *elav-GAL4* driver. Note that the anti-Su(var)3-9 antibody recognizes both endogenous and LacI tagged protein, but not the human SUV3-9. Anti-HA antibody has been used for specific detection of hSUV39H1-HA. Loading control, Vibrator, Vib. (**B**) Western blots showing the expression levels of TBPH in protein extracts obtained from larval brains (right panel) or adult heads at 3 or 20 dpe (left panel) in absence or in presence of chaetocin 100nM. Note that cheatocin treatment induces a reduction of both the heterochromatic markers H3K9me2 and me3 and a significant increase of TBPH levels. Numbers below each blot represent band quantification, normalized on internal loading (actin or Vibrator, Vib). Similar results were obtained in at least 3 biological repetitions. (**C**) Western Blot showing the TBPH protein levels in *G9a* homozygous mutants ($G9a^{RG5}/G9a^{RG5}$) or eggless mutants ($egg^{1473}/egg^{1473}$) compared to wild type controls ($w^{1118}$) in young (3 dpe) fly head extracts. *eggless* mutant flies ($egg^{1473}/egg^{1473}$) die at 3 dpe and are really sick. Numbers below represent band quantification normalized on internal loading (Vibrator, Vib; average of 4 experiments). (**D**) Western Blot showing the TBPH protein levels in *G9a* homozygous mutants ($G9a^{RG5}/G9a^{RG5}$) compared to wild type controls ($w^{1118}$) in young (3dpe) or old (20dpe) fly head extracts. Numbers below represent band quantification normalized on internal loading (Vibrator, Vib; average of 4 experiments). (**E**) Climbing assay performed in *G9a* homozygous mutant ($G9a^{RG5}/ G9a^{RG5}$) and in control flies ($w^{1118}$), at 3 or 20 dpe. Each dot represents the percentage of flies that reach the top of a 50 ml tube in 10 seconds after being tapped to the bottom. $n \geq 20$ animals for each genotype, at least 5 technical replicates. ****$p<0.000$, ns, not significant, calculated by one-way ANOVA. Error bars represent SEM. (**F**) qRT-PCR showing *dFus/cabeza* mRNA levels in young (3 dpe; full circles) or old (20 dpe; empty-dotted circles) fly head extracts. Error bars represent SEM of three independent experiments ($n = 3$; 3 biological replicates and 3 technical replicates). ns: not significant; Mann-Whitney t-test. (**G**) qRT-PCR showing *Su(var)3-9* mRNA levels in young (3 dpe; full circles) or old (20 dpe; empty-dotted circles) fly head extracts. Error bars represent SEM of four independent experiments ($n = 3$; 3 biological replicates and 3 technical replicates). * $p<0.05$; Mann-Whitney t-test.

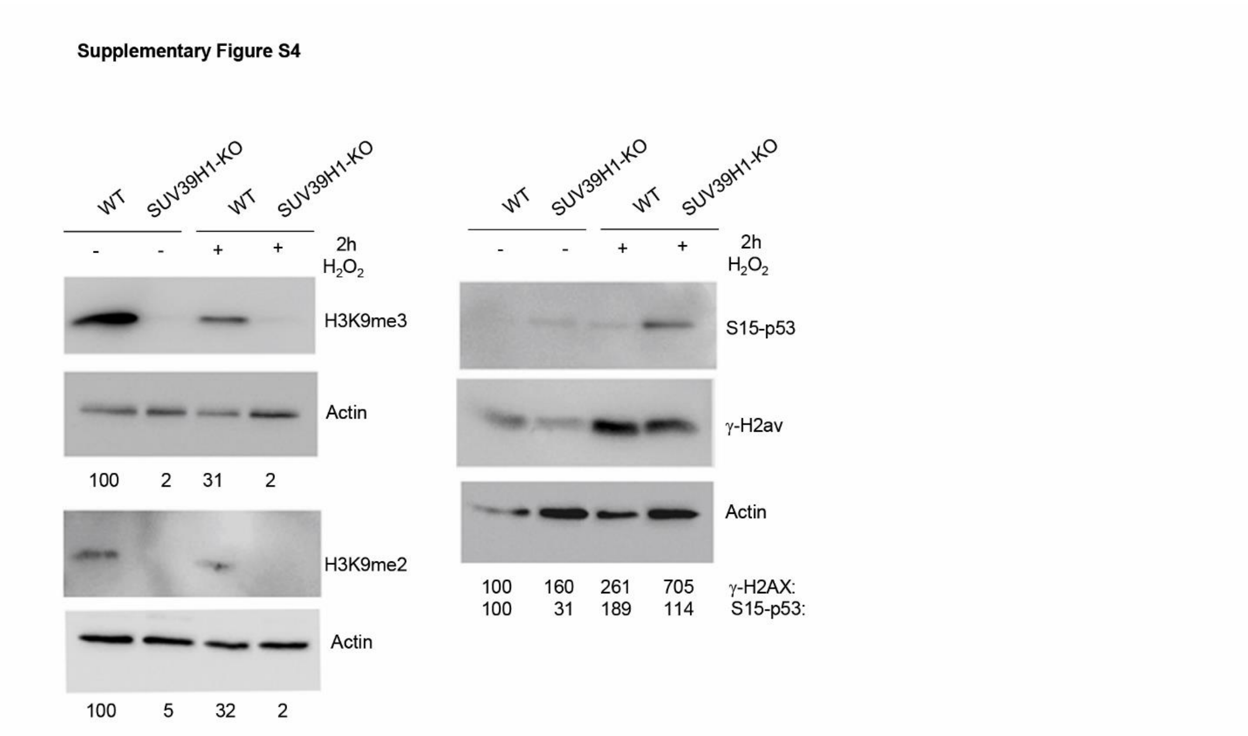


**Supplementary Figure S4. HaCaT human cells exhibit age-related molecular signatures upon $H_2O_2$ treatment**

Western blots showing the expression levels of different aging markers (on the right of each blot) in wild type (WT) or *SUV39H1 KO* HaCaT Keratinocytes after (+) or not (-) treatment with $H_2O_2$ (200mM) for 2 hours (2h). Note that $H_2O_2$ treatment induces a reduction of both the heterochromatic markers H3K9me2 and me3 and an increase of p53-S15 phosphorylated and γ-H2A.X (positive controls of the treatment). Numbers below each blot represent band quantification, normalized on internal loading (Actin). Similar results were obtained in at least 3 biological repetitions.

**Supplementary video V1**

Representative video showing a climbing assay performed on *$w^{1118}$* at 3 dpe (first tube from left), *$w^{1118}$* 20 dpe (middle tube) and *Su(var)3-9* $^{6/1}$ 20 dpe (last tube) flies. The percentage of flies able to reach the top of the 50 ml tube in 10 seconds after being tapped to the bottom is scored to quantify locomotion activity. An average of 15 flies is present in each tube.

### Full Blots

**Full blot**

Figure 1 b

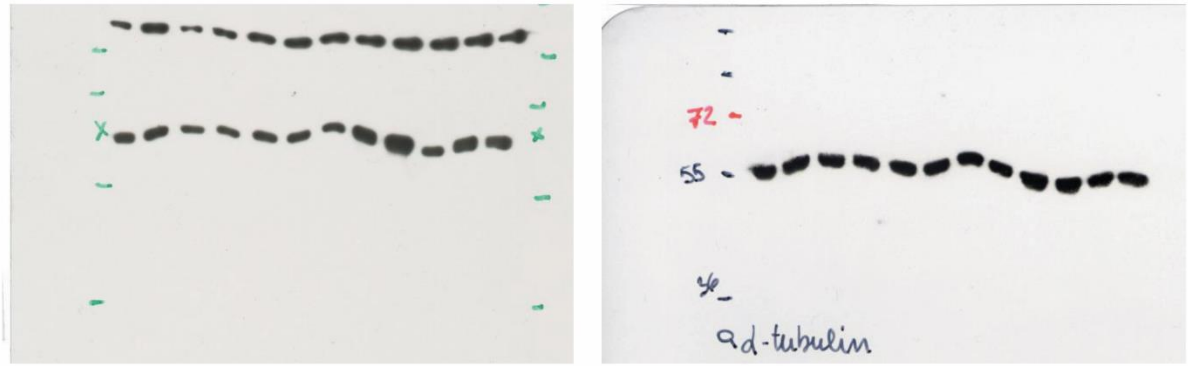


Supplementary Figure S1 b

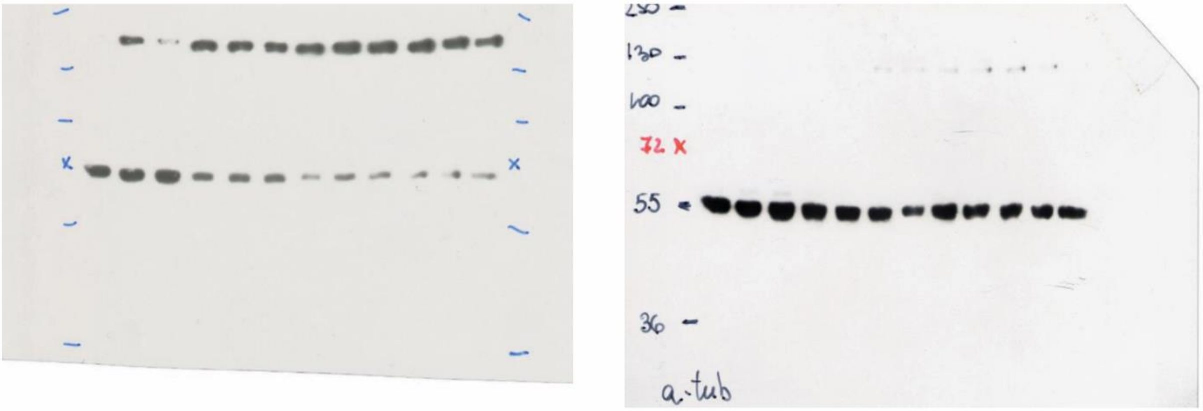

Supplementary Figure 2

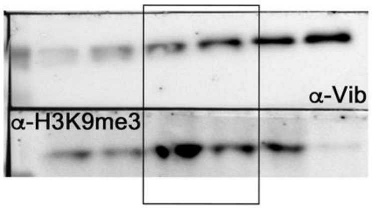


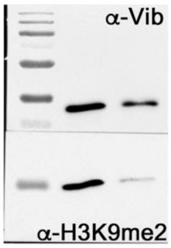


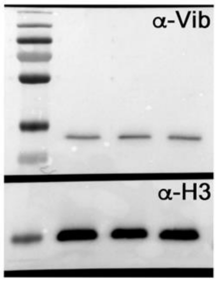


Figure 2

C

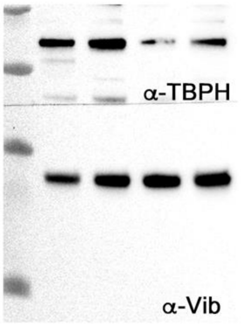


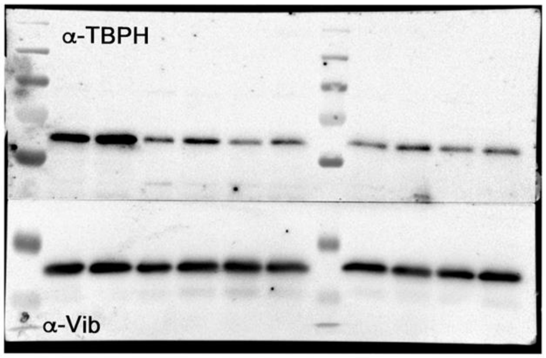


F

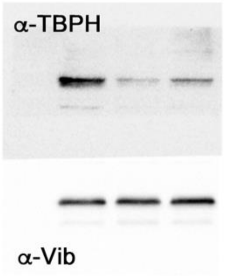

Supplementary Figure 3

A

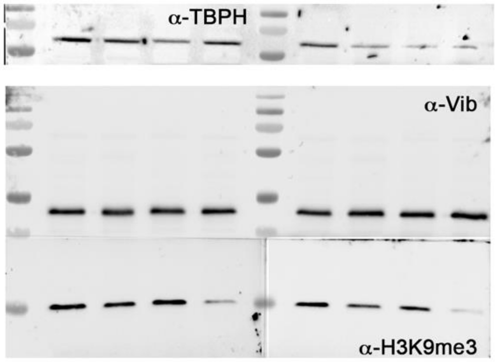


B

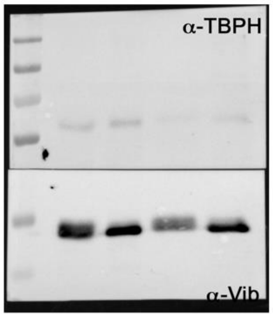


C

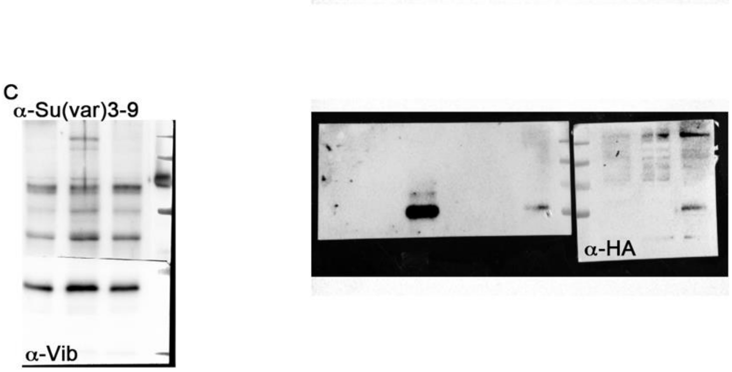


G

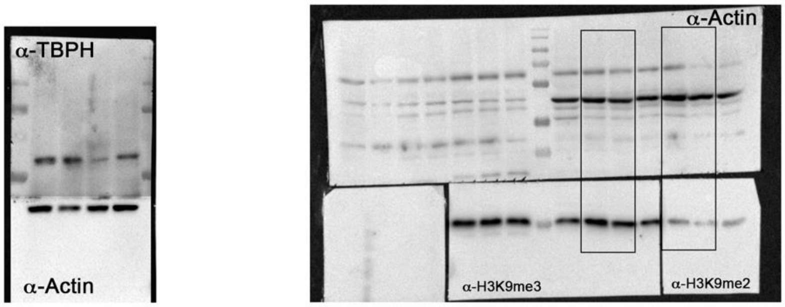


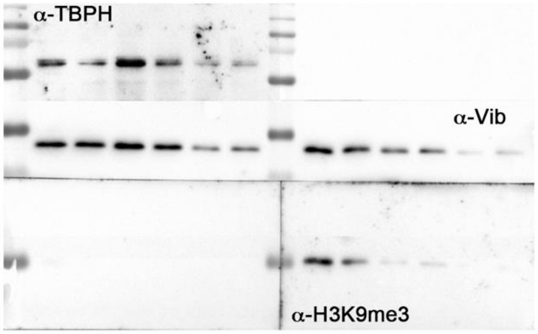

Figure 4A

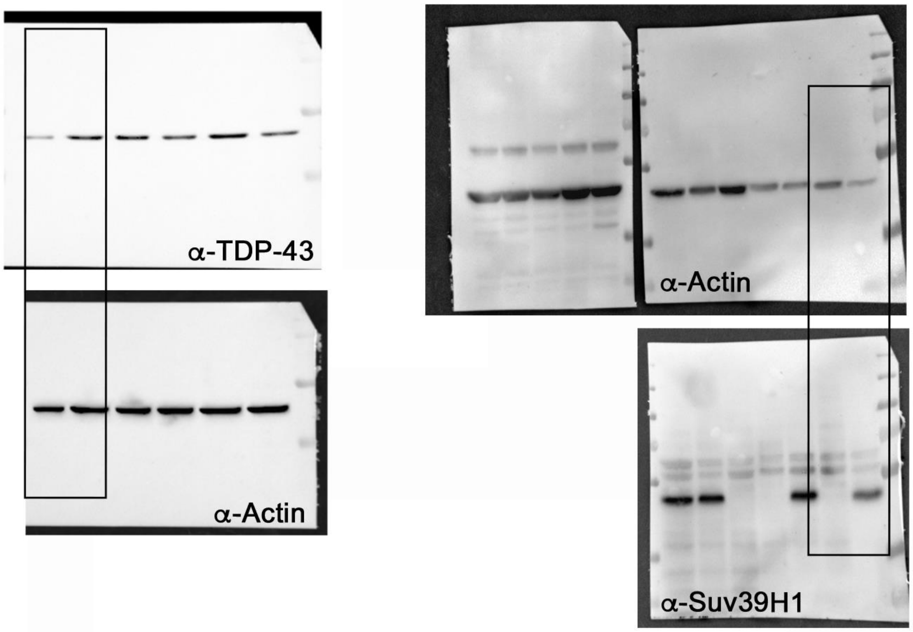


Supplementary Figure 4

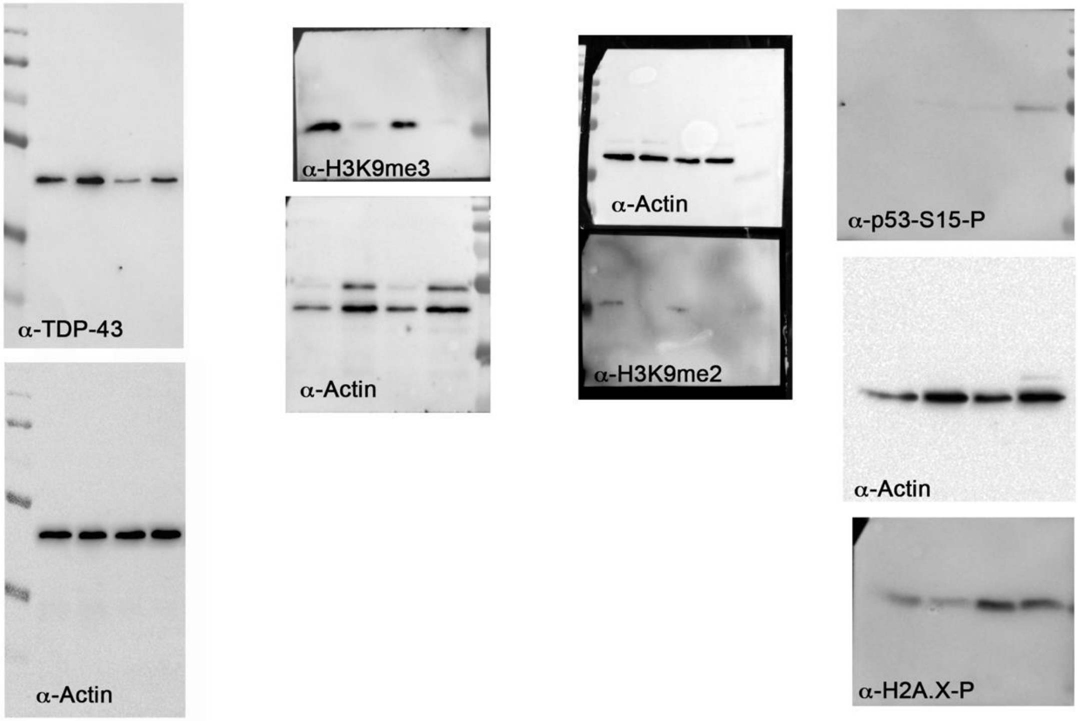
